# Whole-brain afferent input mapping to functionally distinct brainstem noradrenaline cell types

**DOI:** 10.1101/2022.11.22.517460

**Authors:** Jessica Sulkes Cuevas, Mayumi Watanabe, Akira Uematsu, Joshua P. Johansen

## Abstract

The locus coeruleus (LC) is a small region in the pons and the main source of noradrenaline (NA) to the forebrain. While traditional models suggested that all LC-NA neurons project indiscriminately throughout the brain, accumulating evidence indicates that these cells can be heterogeneous based on their anatomical connectivity and behavioral functionality and exhibit distinct coding modes. How LC-NA neuronal subpopulations are endowed with unique functional properties is unclear. Here, we used a viral-genetic approach for mapping anatomical connectivity at different levels of organization based on inputs and outputs of defined cell classes. Specifically, we studied the whole-brain afferent inputs onto two functionally distinct LC-NA neuronal subpopulations which project to amygdala or medial prefrontal cortex (mPFC). We found that the global input distribution is similar for both LC-NA neuronal subpopulations. However, finer analysis demonstrated important differences in inputs from specific brain regions. Moreover, sex related differences were apparent, but only in inputs to amygdala-projecting LC-NA neurons. These findings reveal a cell type and sex specific afferent input organization which could allow for context dependent and target specific control of NA outflow to forebrain structures involved in emotional control and decision making.

## Introduction

The brainstem locus coeruleus (LC) provides most of the noradrenaline (NA) innervation of the forebrain from a small population cells (~2,300 neurons in mice) (Berger et al., 1979; Touret et al., 1982; for recent reviews, see Uematsu et al., 2015; Schwarz & Luo, 2015; Poe et al., 2020; Breton-Provencher et al., 2021) and regulates diverse behavioral functions. Traditionally, the LC was considered a homogeneous region sending broadly collateralized output to many brain regions (Nagai et al., 1981; Room et al., 1981; Schwarz et al., 2015; Kim et al., 2016). However, recent studies reported that some subsets of LC neurons exhibit more specific efferent connectivity (Simpson et al., 1997; Chandler et al., 2014; Kebschull et al., 2016; Hirschberg et al., 2017; Uematsu et al., 2017; Plummer et al., 2020) while others project broadly throughout the brain (Simpson et al., 1997; Steindler, 1981; Schwarz et al., 2015; Kebschull et al., 2016). Furthermore, subpopulations of LC neurons are activated by task relevant stimuli or form distinct cell assemblies (Aston-Jones et al., 1999; Bouret & Richmond, 2009, 2015; Kalwani et al., 2014, Totah et al., 2018). Notably, distinct populations of projection-defined LC-NA neurons exhibit context dependent modular coding (Likhtik & Johansen, 2019; Poe et al., 2020), responding as an ensemble in certain instances or individually in others (Uematsu et al., 2017). Supporting this, a recent study reported that activity in LC-noradrenaline outputs to distinct target regions encode distinct features of a sensory decision making task (Breton-Provencher et al., 2022). In addition, specific populations of projection-defined neurons serve unique behavioral functions. For example, two distinct LC subpopulations project preferentially to the amygdala or the medial prefrontal cortex (mPFC) and facilitate aversive associative learning and anxiety-like behaviors or extinction of aversive emotional memories and exploration, respectively (Uematsu et al., 2015; Borodovitsyna et al., 2020). Despite this accumulating evidence of cellular heterogeneity in the NA system, the mechanisms which imbue distinct LC-NA cell populations with their functional specificity while also allowing them to cooperate as a global ensemble with other neurons in certain contexts is not known.

One potential mechanism for enabling context dependent modular coding and distinct behavioral functionality is through a diversity of afferent input organization, with some inputs to functionally distinct LC-NA cell populations being specific while others being shared across different populations. Prior studies examining inputs to anatomically/genetically, but not functionally, defined cell populations found largely shared inputs across cell groups (Schwarz et al., 2015). However, it is not clear if this is also the case for distinct, functionally defined subpopulations of LC neurons.

To test this hypothesis, we used trans-synaptic rabies virus tracing to examine the brain-wide afferent inputs to either amygdala- or mPFC-projecting LC-NA subpopulations in male and female mice. We took advantage of the hierarchical brain region classification scheme in the Allen Mouse Brain Reference Atlas to determine the afferent input organization at gross, moderate and fine levels of analysis onto these different cell populations. Because the LC is sexually dimorphic and LC-NA neurons as a whole exhibit sex-related differences in afferent input patterns (Sun et al., 2020), we also examined sex differences in input connectivity. Our findings reveal that both cell populations receive inputs from many of the same brain regions, particularly at gross levels of organization. However, at the level of specific brain regions, these cell groups receive a differentiated pattern of inputs which is partially related to sex differences.

## Results

### LC projections to the amygdala and medial prefrontal cortex in the mouse

As initial studies of amygdala- and mPFC-projecting subpopulations were conducted in rats, we first assessed the existence of these subpopulations in the mouse LC. To do this, two different colored retrograde tracers (RetroBeads) were injected into the lateral and basal amygdala (LA/B) and the infralimbic cortex (IL), a subregion of the medial prefrontal cortex (mPFC), of wild-type mice (Fig. 1a-c). Similar to rat, LC-NA projections to IL-mPFC and LA/B were mostly specific as manifested in a small percentage of overlapping retrolabeled cells (from total projecting cells counted, 64% project preferentially to the mPFC, 28% to the amygdala and 7% to both, female and male combined, *n* = 4) (Fig. 1c). However, female mice showed a higher overlap than male mice (12% in females versus 2% in males, *n* = 2 for females and males) (Supplementary Fig. 1). As in the rat, these LC neuronal subpopulations were intermixed and not topographically organized (data not shown).

**Figure 1.**
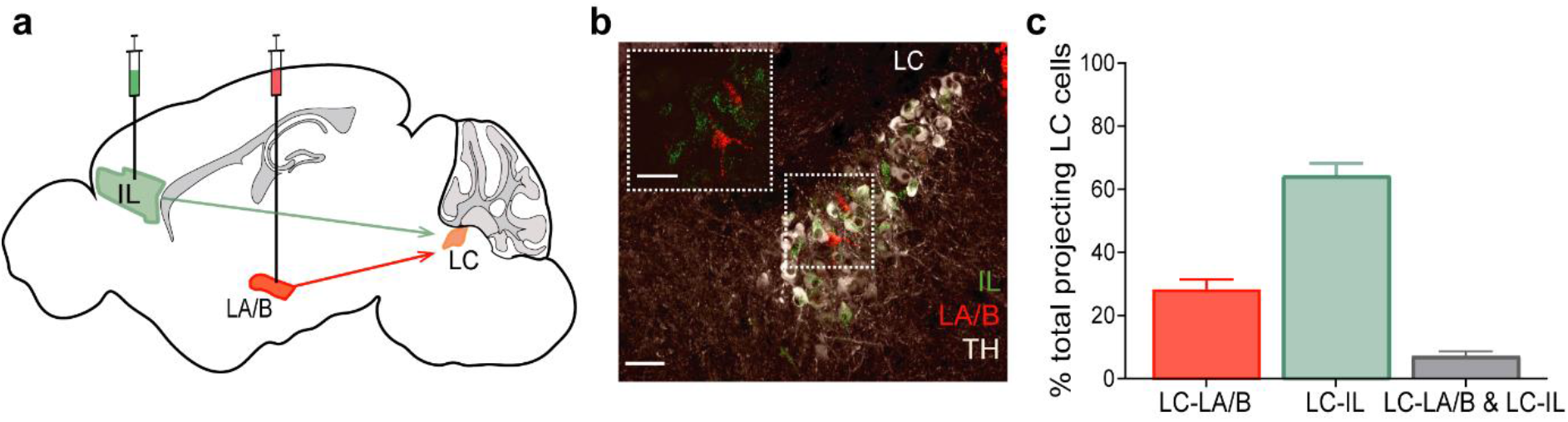
Locus coeruleus (LC) projections to lateral and basal nuclei of amygdala (LA/B) and infralimbic cortex (IL). **a)** RetroBead injections into LA/B and IL. **b)** Example image of male LC: LC-NA cells express tyrosine hydroxylase (TH, white). Inset: red RetroBead-expressing neurons projecting to LA/B and green expressing neurons project to IL. Scale: 50 μm. **c)** Percentage of total LC-NA cells projecting to LA/B, IL or both output regions (n= 4, mean, s.e.m.).

### Whole-brain input distribution to LC subpopulations in the mice

#### Starter cells and total input distribution

To evaluate the whole-brain input distribution of LC neurons projecting to amygdala or mPFC, we used a retrograde rabies-mediated cTRIO labeling approach (“cell-type-specific tracing the relationship between input and output”, Schwarz et al., 2015). In mice expressing Cre recombinase in noradrenergic neurons (noradrenaline transporter-Cre, NAT-cre), we injected a combination of retrograde viruses (canine adenovirus, CAV and retro-adeno-associated virus, AAV) expressing cre-dependent Flp recombinase in amygdala or mPFC (CAV-FLEx-Flp and retrograde AAV-FLEx-flpO, combined to avoid reduce potential viral tropism effects) and Flp-dependent rabies helper viruses (AAV-FLEx(FRT)-TVA-mCherry and AAV-fDIO-H2B-3xHA-N2cG) in the LC (Fig. 2a). This combination of cre- and flp-dependent viral delivery allowed specific expression of the helper viruses in LC-NA neurons projecting to amygdala or mPFC. Three to four weeks later, rabies virus was injected into the LC (EnVA-CVS-N2cΔG-EGFP) (Schwarz et al., 2015; Soya et al., 2017).

Starter cells were defined as LC-noradrenaline neurons (tyrosine hydroxylase (TH)-expressing cells, immunolabeled with Alexa Fluor 647) co-expressing a nuclear localized (fused to histone H2B) HA tag (label for glycoprotein, immunolabeled with Alexa Fluor 594, see Methods), mCherry in cytoplasm (TVA receptor) and EGFP (rabies). All starter cells (i.e., those expressing nuclear HA-tag, cytoplasmic mCherry and EGFP) were TH positive, and no co-expression of helper viruses and rabies virus was found outside LC-noradrenaline neurons (Fig. 2a, b). Both amygdala- and mPFC-projecting LC subpopulations showed similar numbers of starter cells (Fig. 2c, amygdala-projecting starter cells: 13 ± 2 neurons versus mPFC-projecting starter cells: 10 ± 2 neurons, mean ± s.e.m.). Quantification of input cells showed that amygdala-projecting LC neurons received double the number of input cells (Fig. 2d, inputs to amygdala-projecting cells: 4485 ± 834 neurons versus inputs to mPFC-projecting cells: 2168 ± 402 neurons). The convergence index (the ratio of all inputs to all starter cells for each group) showed that both subpopulations received similar number of total inputs from the whole-brain (Fig. 2e, 407 ± 101 neurons for the amygdala-projecting group, 246 ± 56 neurons for the mPFC-projecting group. Mann-Whitney U= 17, no significant differences; median_LC-amygdala_= 211.8, *n* = 7; median_LC-mPFC_= 273.4, *n* = 8).

**Figure 2.**
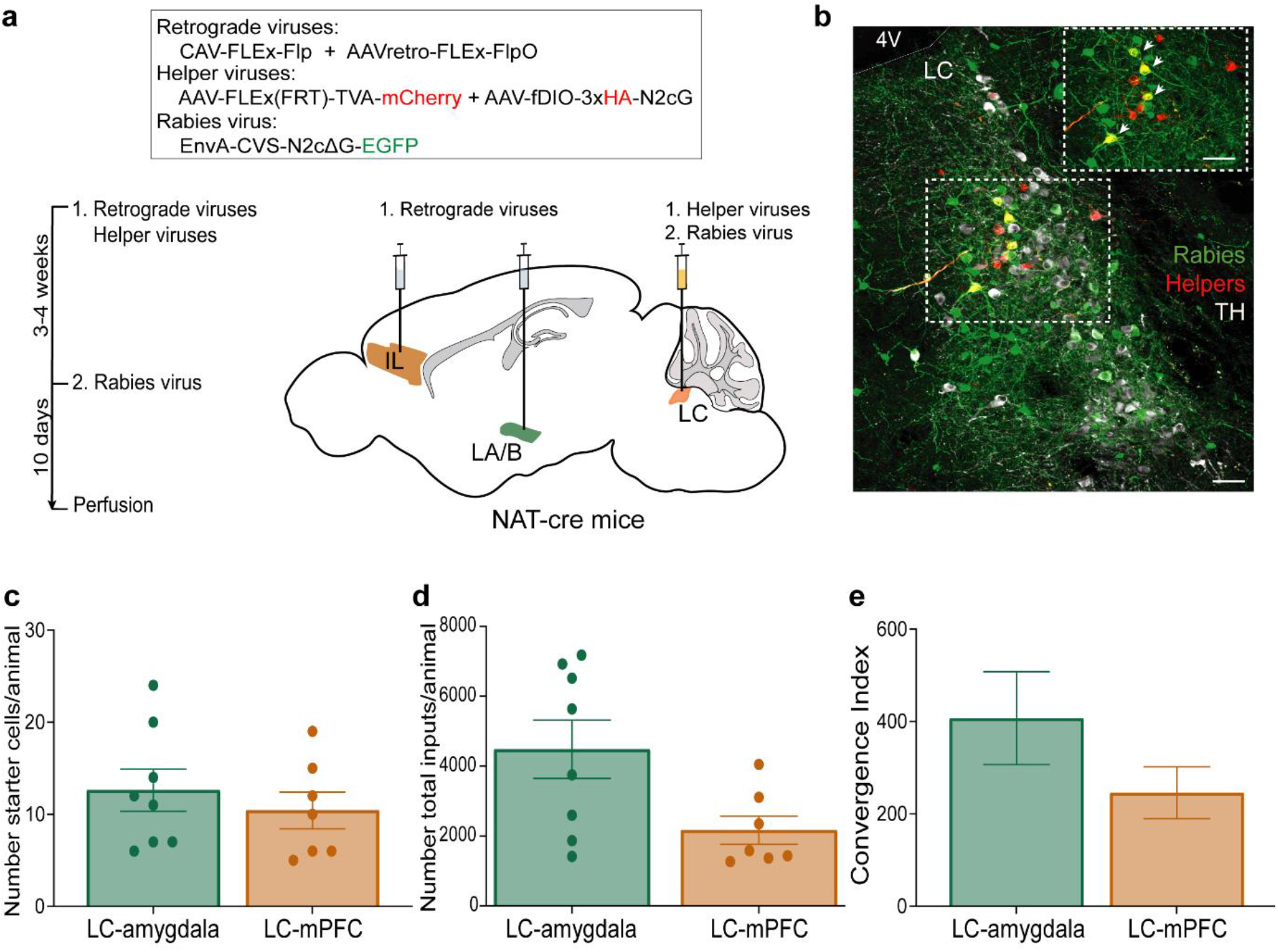
Cell-type specific trans-synaptic rabies viral input mapping approach. **a**) Top: viruses used for trans-synaptic rabies expression. Bottom left: timeline of virus injections. Bottom right: injection sites. **b**) Example image of starter cells in LC. Inset: close-up image of starter cells denoted by white arrowheads. White cells: TH-expressing cells. Red cells: helper virus-expressing cells. Green cells: rabies virus-expressing cells. 4V: fourth ventricle. Scale: 50 μm. **c**) Number of starter cells for amygdala-projecting (LC-amygdala) and mPFC-projecting (LC-mPFC) LC-NA neurons. **d**) Number of total inputs for LC-amygdala and LC-mPFC neuronal populations. **e**) Convergence Index for LC-amygdala and LC-mPFC cell groups (LC-amygdala, n= 7; LC-mPFC, n= 8; mean ± s.e.m.). Circles represent individual animal values.

#### Whole-brain input analysis workflow

To develop the whole-brain atlas of inputs to amygdala- and mPFC-projecting LC cells, 50-μm coronal sections were sampled from every 3^rd^ section throughout the entire brain (approximately from bregma: +3.0 to −7.2), excluding approximately 400 nm around the injection site in LC. Each region was registered and analyzed using the anatomical classification criterion of the Allen Mouse Brain Reference Atlas (Fig. 3). This classification allows for quantification of input distribution in a hierarchical manner (Sunkin et al., 2013; Wang et al., 2020). Specifically, three levels of organization have been proposed (Sun et al., 2020): gross, moderate, and fine corresponding to the hierarchical category of anatomical regions according to the Allen Mouse Brain Atlas (see Methods). We used these levels to guide our quantification of inputs to amygdala- and mPFC-projecting LC-NA neurons.

**Figure 3.**
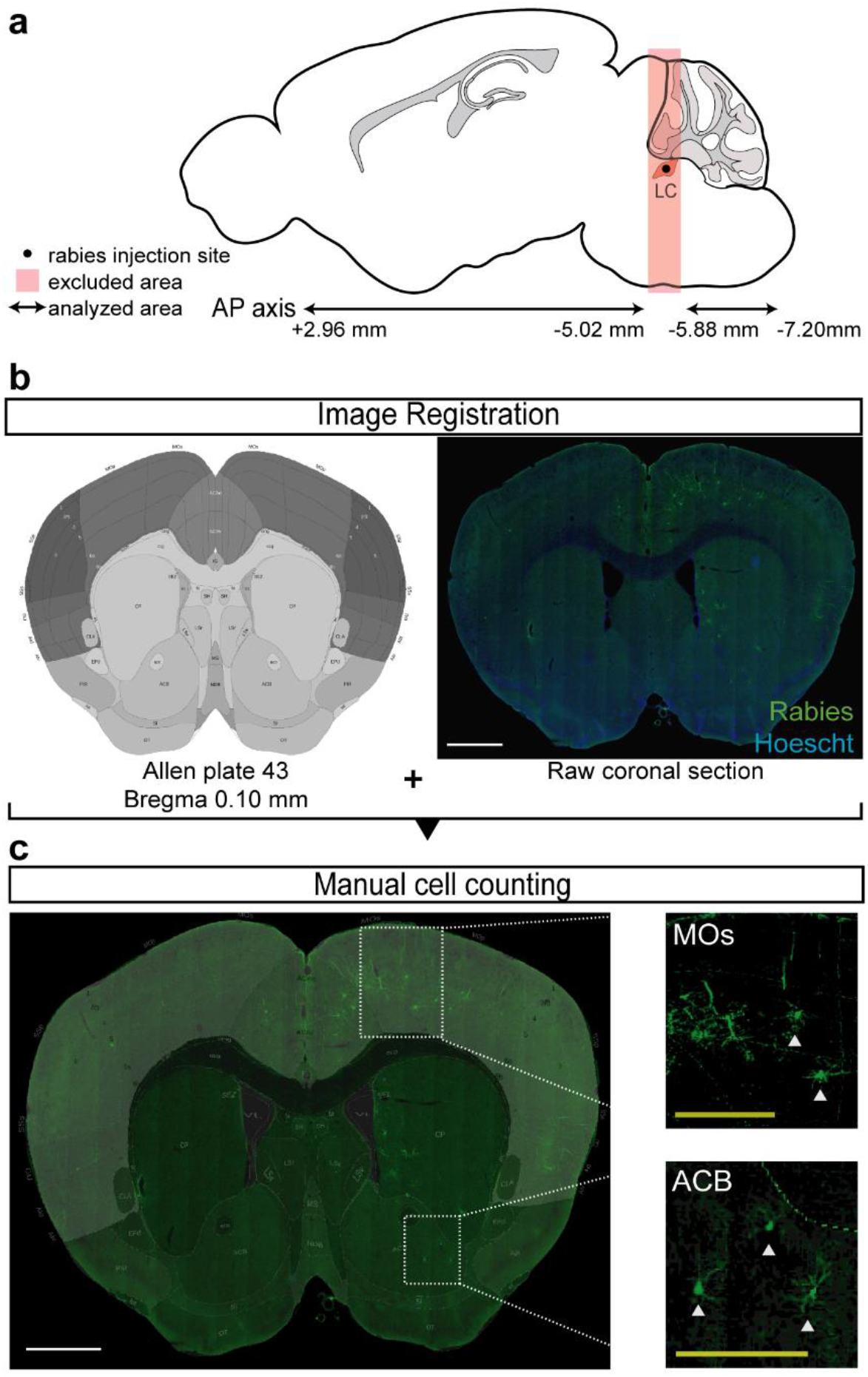
Whole-brain input distribution analysis workflow. **a**) Schematic analyzed. (See Methods). AP axis: anterior-posterior axis. **b**) Image registration. templates from Allen Mouse Brain Reference Atlas were overlaid to coronal sections (blue: Hoechst dye; green: rabies) **c**) Graphical depiction of coronal sections in which cells were manually count. Scale, white: 200 μm. Scale, yellow: 100 μm. MOs: motor areas, secondary; ACB: nucleus accumbens.

### Gross level of organization

To visualize the whole-brain input distribution at the gross level of analysis, the number of input neurons to amygdala- or mPFC-projecting LC cells was quantified in each broad-category anatomical input area. Input proportions from these areas were quantified to determine the relative strength of inputs to the distinct LC neuronal subpopulations (Fig. 4).

**Figure 4.**
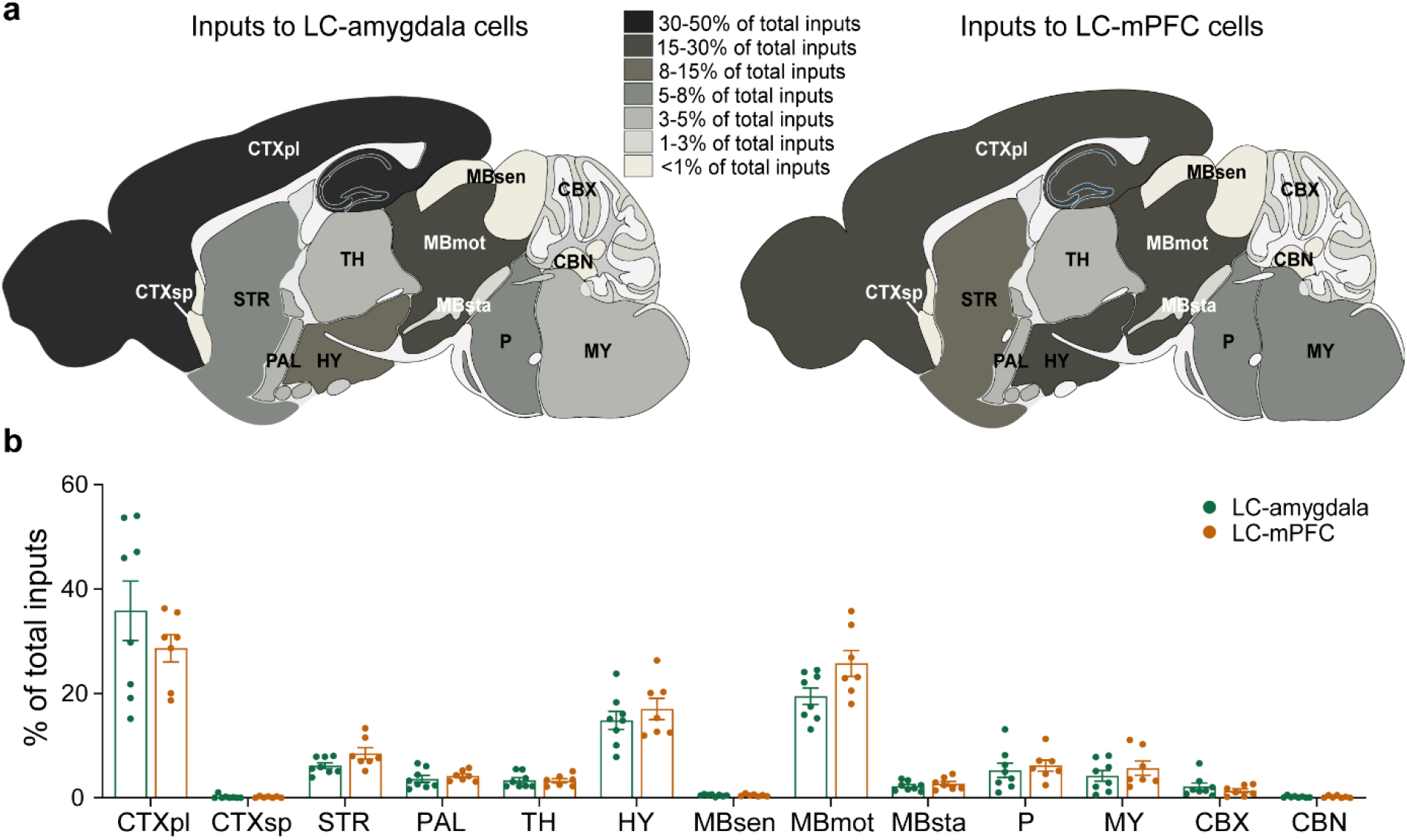
Whole-brain inputs to LC-NA subpopulations at the gross level of organization. **a)** Whole-brain input distribution. Left: proportion of inputs to amygdala-projecting LC cells (LC-amygdala).Right: proportion of inputs to mPFC-projecting LC cells (LC-mPFC). **b)** Percentage of total inputs from the whole-brain to LC-amygdala and LC-mPFC cell populations (LC-amygdala, n= 7; LC-mPFC, n= 8, mean ± s.e.m.). Circles represent individual animal values. CTXpl: cortex, plate. CTXsp: cortex, subplate. STR: striatum. PAL: pallidum. TH: thalamus. HY: hypothalamus. MBsen: midbrain, sensory-related. MBmot: midbrain, motor related. MBsta: midbrain, behavioral state related. P: pons. MY: medulla. CBX: cerebellar cortex. CBN: cerebellar nuclei.

Cortex plate, hypothalamus and motor-related midbrain were the main source of inputs to both subpopulations. The distribution of inputs from all areas was largely similar with no apparent statistically significant differences (Fig. 4a). However, qualitatively, amygdala-projecting LC neurons received more inputs from the cortex plate than mPFC-projecting cells (35.8 ± 5.7 % versus 28.7 ± 2.6 %, respectively) while mPFC-projecting cells received more input from the hypothalamus (to amygdala-projecting LC neurons: 14.9 ± 1.7 %, to mPFC-projecting LC neurons: 17 ± 2 %) and the motor-related midbrain (MBmot) areas (to amygdala-projecting LC neurons: 19.5 ± 1.6 %, to mPFC-projecting LC neurons: 25.8 ± 2.5) (Fig. 4b)

### Moderate level of organization

We next examined lower hierarchical levels of organization to determine whether individual brain regions provide more specific synaptic input to these distinct LC-noradrenaline cell populations. For moderate and fine levels of organization, input regions were defined as those representing ≥ 0.1 % of the total fraction of inputs (Sun et al., 2020). Moderate level of organization corresponds mainly to subregions of the 5^th^ “node” of the Allen Mouse Brain Atlas categorization (Fig. 5a). While the density of many inputs to the different LC cell subpopulations at this level was similar, mPFC-projecting LC neurons received significantly more inputs from the substantia nigra, compact (SNc, U= 7, P= 0.014, median_LC-amygdala_= 0.00085; median_LC-mPFC_= 0.00057, Fig. 5b, d) and the red nucleus (RN, U= 10, P= 0.04, median_LC-amygdala_= 0.00996; median_LC-mPFC_= 0.00318, Fig. 5c, e), both regions within the motor-related midbrain broad category.

**Figure 5.**
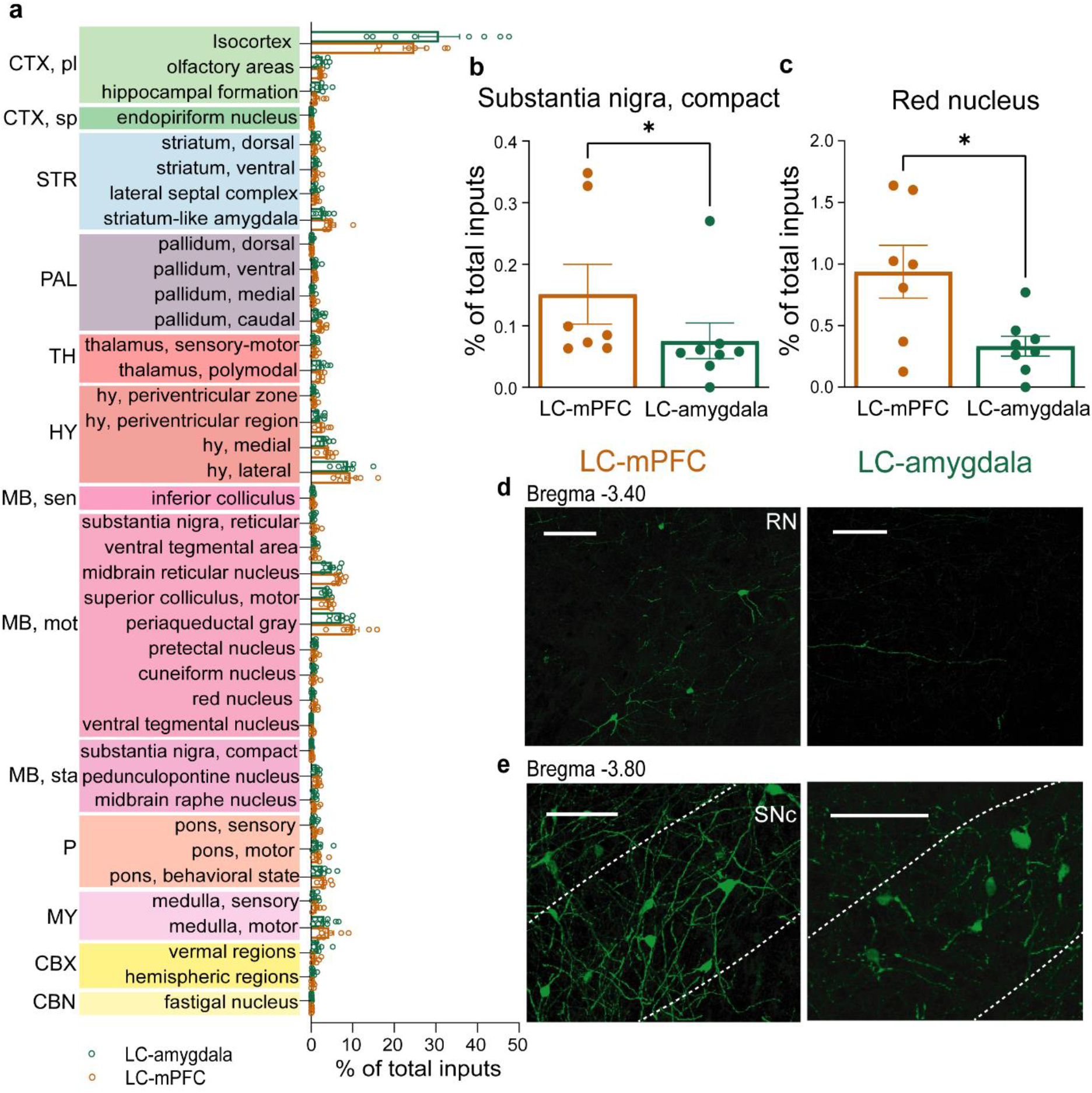
Whole-brain inputs to LC-NA subpopulations at the moderate level of organization. **a**) Percentage of total inputs at the moderate level of organization. **b**) Percentage of total inputs to LC-mPFC and LC-amygdala subpopulations from the red nucleus (RN). **c**) Percentage of total inputs to LC-mPFC and LC-amygdala projecting subpopulations from the substantia nigra, compact (SNc). * P < 0.05. Circles represent individual animal values. **d-e**) Example images of input cells in RN and SNc to LC-mPFC (left) and LC-amygdala (right). Scale: 100 μm

### Fine level of organization

Finally, we evaluated the distribution of inputs from specific brain areas at the fine level of organization (related to the 6^th^-7^th^ “nodes” of the Allen Mouse Brain Atlas). This level contains specific subregions and subnuclei within the broader regional groupings and includes around 80 individual presynaptic input areas to LC subpopulations.

The amygdala-projecting LC-NA neurons received more inputs than mPFC-projecting LC cells from a variety of regions. These include specific nuclei within the isocortex including the visual areas (VIS, U= 7, P= 0.013, median_LC-amygdala_= 0.00139; median_LC-mPFC_= 0.00481, Fig. 6a, b) and the posterior parietal association areas (PTLp, U= 9, P= 0.029, median_LC-amygdala_= 0.00139; median_LC-mPFC_= 0.00377, Fig. 6c, d). From the olfactory areas, the piriform area also projected more strongly to this LC-NA cell population (PIR, U= 6, P= 0.009, median_LC-amygdala_= 0.00425; median_LC-mPFC_= 0.01244, Fig. 6e, f). Finally, amygdala-projecting LC cells also received denser innervation from the fundus of striatum (FS, U= 10, P= 0.039, median_LC-amygdala_= 0.00064; median_LC-mPFC_= 0.00136, Fig. 6g, h) and the cerebellar vermal region, the nodulus (NOD, U= 10, P= 0.04, median_LC-amygdala_= 0.00079; median_LC-mPFC_= 0.00201, Fig. 6i, j).

**Figure 6.**
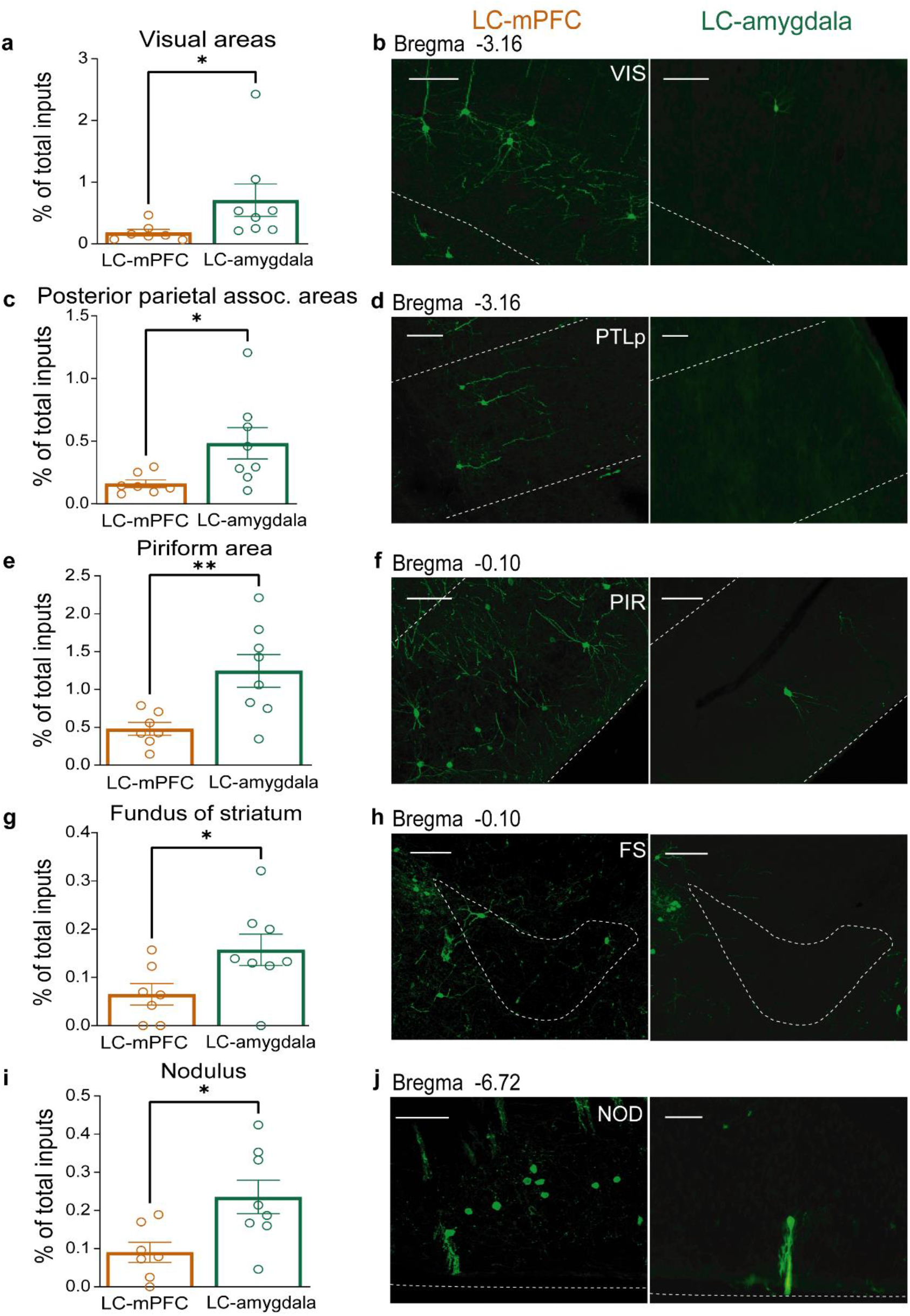
Inputs to amygdala projecting LC-NA neurons at the fine level of organization. Percentage of inputs from specific areas of the isocortex to LC-amygdala and LC-mPFC subpopulations **(a-d)**. **a**) Inputs from the visual areas (VIS). **b**) Example images of inputs from VIS to LC-amygdala (left) and to LC-mPFC (right). **c**) Inputs from the posterior parietal association areas (PTLp). **d**) Example images of inputs from PTLp to LC-amygdala (left) and to LC-mPFC (right). **e**) Percentage of inputs from the olfactory area, piriform area (PIR). **f**) example images of inputs from PIR to LC-amygdala (left) and LC-mPFC (right). **g**) Percentage of inputs from the striatal region, fundus of striatum (FS) to LC-amygdala and LC-mPFC subpopulations. **h**) Example images of inputs from FS to LC-amygdala (left) or LC-mPFC (right). **i**) Percentage of inputs from the cerebellar cortex subregion, nodulus (NOD) to LC-amygdala and LC-mPFC subpopulations.) Example images of inputs from NOD to LC-amygdala (left) or LC-mPFC (right). *P < 0.05, **P < 0.01. Circles represent individual animal values. Scale: 100 μm.

By contrast, mPFC-projecting LC-NA neurons received preferential inputs from a number of subcortical regions. From olfactory areas, the anterior olfactory nucleus (AON) projected more strongly to mPFC-projecting LC-NA neurons (U= 10, P= 0.04, median_LC-amygdala_= 0.01023; median_LC-mPFC_= 0.00297, Fig 7a, b). In the hypothalamus, the parasubthalamic nucleus (PSTN, U= 8, P= 0.0205, median_LC-amygdala_= 0.00691; median_LC-mPFC_= 0.00506, Fig. 7e, f) and the anteroventral periventricular nucleus (AVPV, U= 7, P= 0.014, median_LC-amygdala_= 0.00126; median_LC-mPFC_= 0.0037, Fig. 7g, h) projected preferentially to this cell population. Interestingly, the central nucleus of the amygdala (CeA), categorized as striatum-like amygdala by the Allen Brain Reference Atlas, sent dominant inputs to the mPFC-projecting LC-NA neurons (U= 7, P= 0.014 median_LC-amygdala_= 0.03874, *n* = 7; median_LC-mPFC_= 0.02019, Fig. 7c, d).

**Figure 7.**
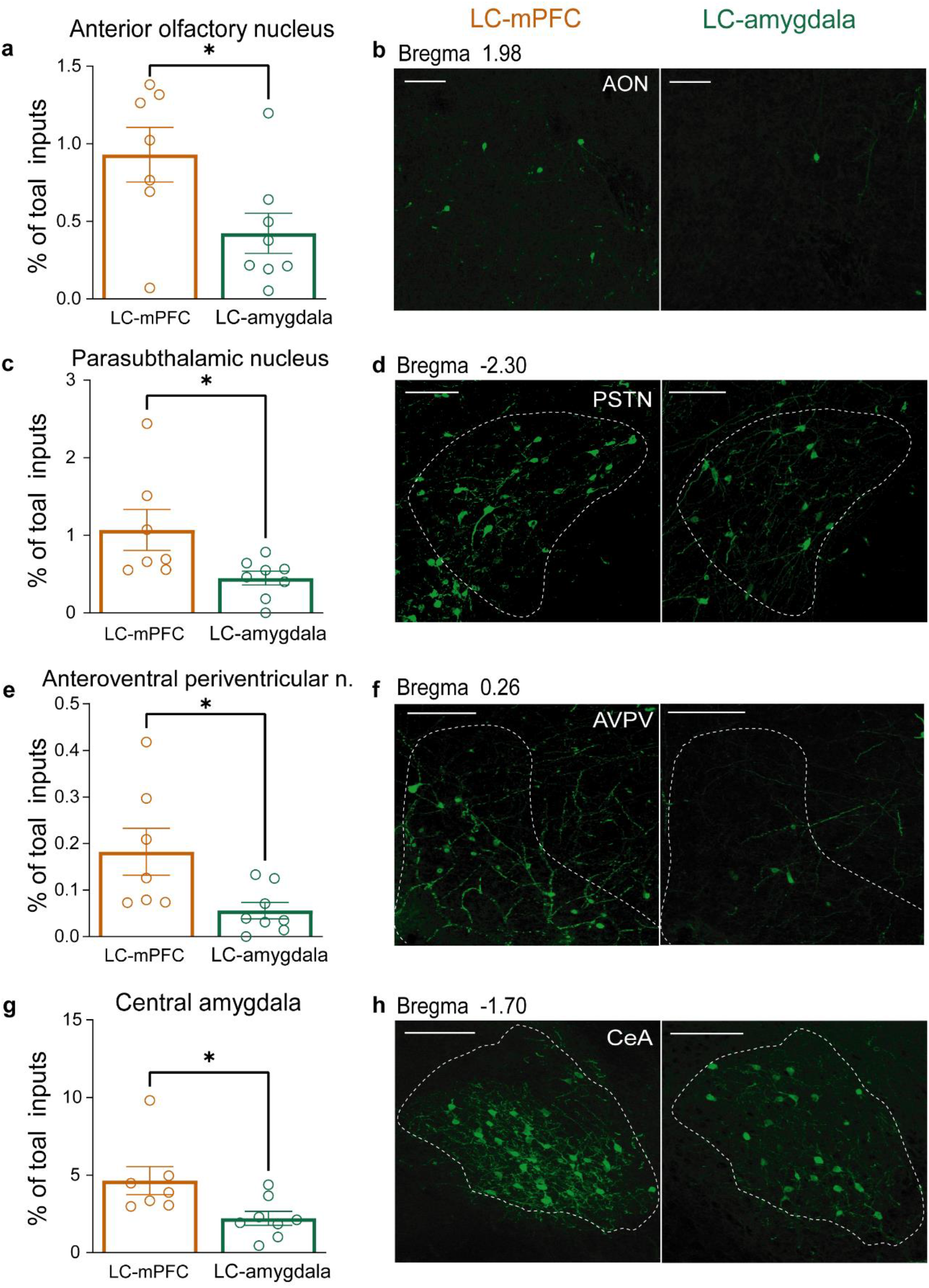
Whole-brain inputs to mPFC projecting neurons at the fine level of organization. **a)** Percentage of inputs from the anterior olfactory nucleus (AON) to LC-mPFC and LC-amygdala subpopulations. **b**) Example images of inputs from AON to LC-mPFC (left) and to LC-amygdala (right) subpopulations. **c-f**) Percentage of inputs from hypothalamic regions to LC-mPFC and LC-amygdala subpopulations: inputs from the parasubthalamic nucleus (PTSN, **c**) with example images to LC-mPFC (left) and to LC-amygdala (right) subpopulations (**d**). Inputs from the anteroventral periventricular nucleus (AVPV, **e**) and example images to LC-mPFC (left) and to LC-amygdala (right) subpopulations (**f**). **g**) Percentage of inputs from the striatum-like region, central amygdala (CeA) to LC-mPFC and LC-amygdala subpopulations. **h**) Example images of inputs from CeA to LC-mPFC (left) and to LC-amygdala (right) subpopulations. *P < 0.05. Circles represent individual animal values. Scale: 100 μm.

The analysis of input distribution patterns at different categorical levels reveals how globally, both LC subpopulations receive similar afferent inputs, consistent with previous results (Schwarz et al., 2015). However, a finer level of analysis, demonstrated that specific regions project preferentially to amygdala- or mPFC-projecting LC-NA cell populations. Together, the combination of shared and specific inputs suggests a mechanism for context dependent global or specific recruitment of these distinct cell populations.

### Sex-related input distribution differentiation

Because of the sexually dimorphic nature of the LC (Pinos et al., 2001; Helena et al., 2009; Bangasser et al., 2011; 2016, 2018; Guajardo et al., 2017; Sun et al., 2020) and the fact that LC-NA neurons receive partially distinct synaptic inputs in female and male animals (Sun et al., 2020), we next assessed sex-related differences in the presynaptic inputs to the different LC-NA neuronal subpopulations.

Inputs to amygdala- and mPFC-projecting LC-NA neurons were evaluated in female and male mice (Supplementary Fig. 2). The number of starter cells in all groups was similar (Supplementary Fig. 2b, LC-amygdala, females: 12 ± 4 cells, *n* = 4; males: 13 ± 3 cells, *n* = 4. LC-mPFC, females: 7 ± 2 cells, *n* = 3; males: 13 ± 3 cells, *n* = 4). The number of input cells for each LC neuronal population was comparable in both female and male animals (Supplementary Fig. 2c, LC-amygdala, females: 5287 ± 1333 cells; males: 3683 ± 1020 cells. LC-mPFC, females: 1973 ± 571 cells; males: 2315 ± 622 cells). Consistent with sex-combined data, a convergence index (the ratio of all inputs to all starter cells for each group) analysis showed that the different LC neuronal subpopulations for each sex received a similar number of inputs per cell (Supplementary Fig. 2d, in amygdala-projecting groups: females: 520 ± 187 neurons, males: 295 ± 63 neurons. U= 5, no significant differences; median_femaleLC-amygdala_= 444, *n* = 4; median_maleLC-amygdala_= 264. In mPFC-projecting groups: females: 314 ± 111 neurons,males:195 ± 53 neurons.U= 3, no significant differences; median_femaleLC-mPFC_= 287, *n* = 3; median_maleLC-mPFC_= 168, *n* = 4).

While no statistical differences were detected in inputs to these cell populations at the gross level of organization, qualitative examination suggested that in females, amygdala-projecting LC-NA neurons tended to receive more inputs from cortex plate, medulla, and cerebellar cortex than male animals. By contrast, this cell population tended to receive less inputs from hypothalamus and pons than males (Supplementary Fig. 3a, b). mPFC-projecting LC neurons in female mice tended to receive more inputs from striatum, hypothalamus, pons, medulla, and cerebellar cortex, but less inputs from cortex plate and thalamus than males (Supplementary Fig. 3c, d).

At moderate and fine levels of organization, sex-related differences in input connectivity were detected, but only to the amygdala-projecting LC-NA subpopulation. Amygdala-projecting LC-NA neurons in female mice showed dominant inputs from the hippocampal region of the cortex plate (Fig. 8a, U= 0, P= 0.029; median_femaleLC-amygdala_= 0.02437; median_maleLC-amygdala_= 0.00587), in particular from the CA3 subregion (Fig. 8b, U= 0, P=0.029; median_femaleLC-amygdala_= 0.01115; median_maleLC-amygdala_= 0.00046). Moreover, the endopiriform area, a region categorized as part of the cortex subplate, showed preferential connectivity with amygdala-projecting LC-NA neurons in females (Fig. 8 c, d, U= 0, P=0.029; median_femaleLC-amygdala_= 0.0008; median_maleLC-amygdala_= 0).

**Figure 8.**
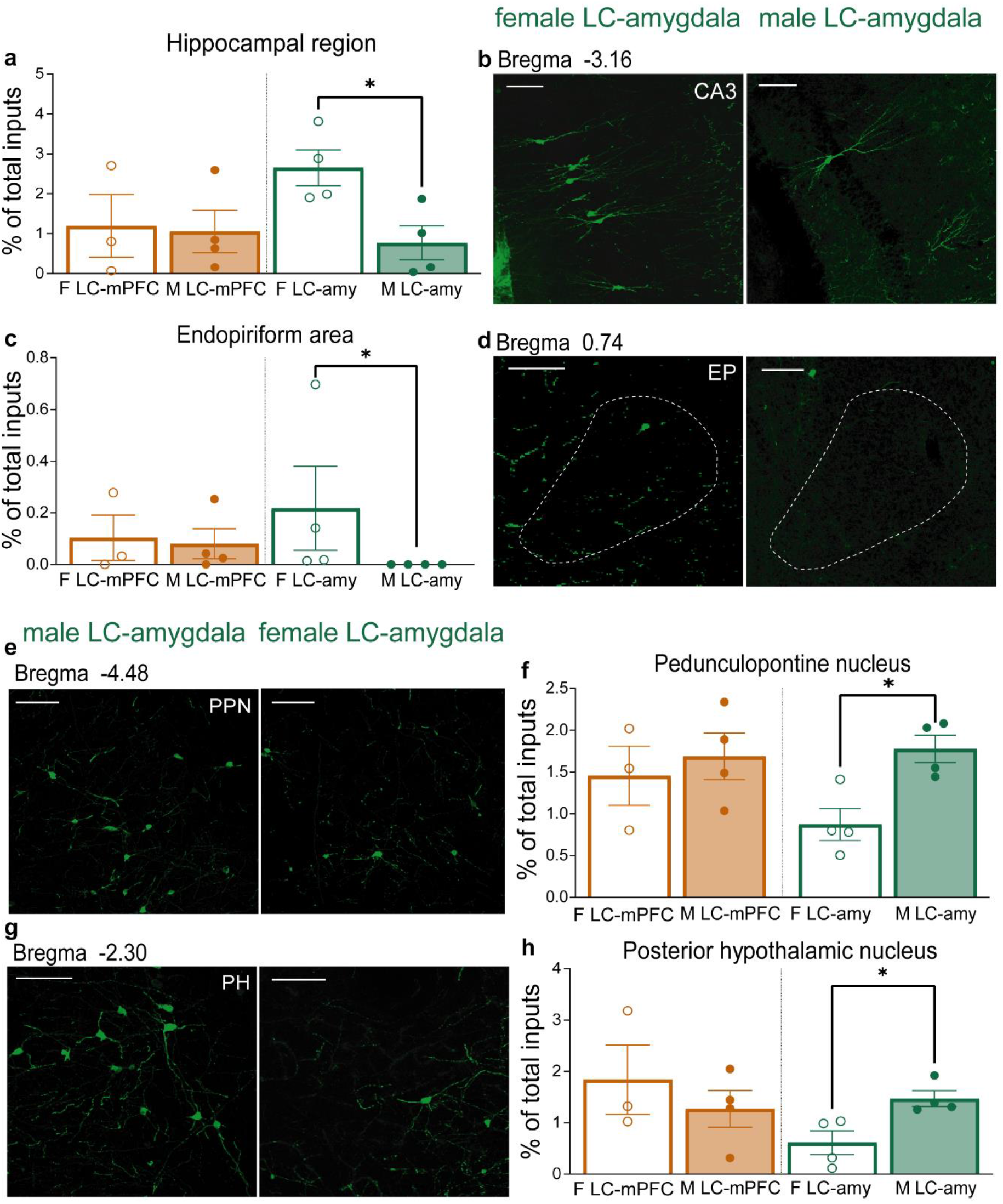
Sex-related differences in input regions to the LC-amygdala subpopulation. Dominant inputs to female LC-amygdala subpopulation **(a-d)**. **a)** Percentage of inputs to female (LC-mPFC, F LC-mPFC; LC-amygdala, F LC-amy) and male (LC-mPFC, M LC-mPFC; LC-amygdala, M LC-amy) from hippocampal region of the isocortex. **b)** Example images of inputs from CA3 of the hippocampal region to F LC-amy (left) and M LC-amy (right). **c)** Percentage of inputs to each subpopulation in female and male mice from the cortex, subplate endopiriform area (EP). **d)** Example images of EP to F LC-amy (left) and M LC-amy (right). Dominant inputs to male LC-amygdala subpopulation **(e-i)**. **e)** Example images from the posterior hypothalamic nucleus (PH) to M LC-amy (left) and F LC-amy (right). **f)** Percentage of inputs to each subpopulation in female and male mice from the PH. **g)** Example images of inputs from the pedunculopontine nucleus (PPN) to M LC-amy (left) and F LC-amy (right). **h)** Percentage of inputs to each subpopulation in female and male mice from the PPN. *P < 0.05. Circles represent individual animal values. Scale: 100 μm.

Input pattern differences onto amygdala-projecting LC-NA neurons were also found in male mice. At the moderate level of organization, differences were apparent in inputs from the sensory-related midbrain pedunculopontine nucleus, which projected strongly to both LC-NA populations, but preferentially to amygdala projecting cells in male compared with female animals (Fig. 8e, f; Supplementary Fig. 4, U= 0, P=0.029; median_maleLC-amygdala_= 0.01789; median_femaleLC-amygdala_= 0.0079). Furthermore, at the fine level of organization, the posterior hypothalamic nucleus projected more to amygdala-projecting LC-NA neurons in male compared to female animals (Fig. 8 g, h, U= 0, P=0.029; median_maleLC-amygdala_= 0.01349; median_femaleLC-amygdala_= 0.00654). These data show that in addition to general differences in inputs to amygdala- and mPFC-projecting LC-NA neurons, sex related differences in inputs onto these specific cell populations also exist.

## Discussion

Emerging evidence indicates that specific populations of LC-NA neurons regulate distinct functions and that, depending on the context or sensory stimulus, LC-NA neurons can be selectively or broadly recruited. However, it is not clear how this distinct functionality and context dependent modular coding arises. Examining the whole-brain inputs to amygdala- and mPFC-projecting LC-NA neurons using a combinatorial molecular and projection cell-type specific trans-synaptic rabies approach (Schwarz et al., 2015; Schwarz & Luo, 2015; Soya et al., 2017) we show that, globally, both subsets of cells receive largely similar afferent inputs. However, a finer level of analysis demonstrated that specific brain structures preferentially target each subpopulation (Fig. 9). Furthermore, inputs onto amygdala-, but not mPFC-projecting LC-NA cells were sexually dimorphic at moderate and fine levels of analysis. These results reveal a partially segregated afferent input architecture onto amygdala- and mPFC-projecting LC-NA neurons, providing a potential circuit basis for the unique behavioral functionality and context dependent activation of these distinct LC cell types. These findings also suggest circuit mechanisms for sexually dimorphic regulation of LC-NA.

**Figure 9.**
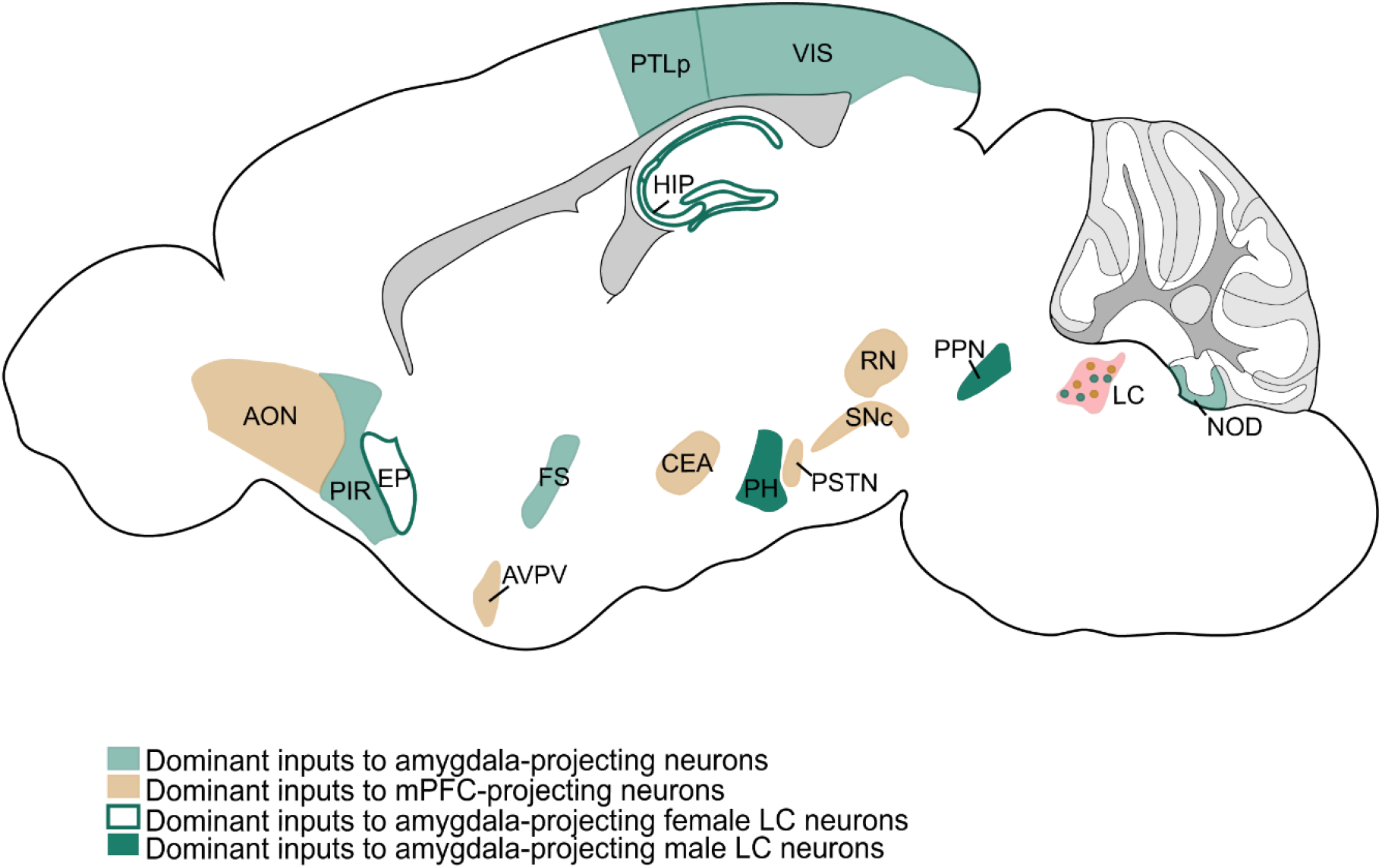
Overview of differences in inputs to amygdala- and mPFC-projecting LC-NA subpopulations. Summary of brain regions which send predominant inputs to either LC-amygdala or LC-mPFC subpopulations and regions that send inputs preferentially to female of male amygdala-projecting LC neurons. AON: anterior olfactory nucleus, AVPV: anteroventral periventricular nucleus, CEA: central amygdala, EP: endopiriform nucleus, FS: fundus of striatum, HIP: hippocampal region, LC: locus coeruleus, NOD: nodulus, PH: posterior hypothalamic nucleus, PIR: piriform area, PPN: pedunculopontine nucleus, PSTN, parasubthalamic nucleus, PTLp: posterior parietal association areas, RN: red nucleus, SNc: substantia nigra, compact, VIS: visual areas.

Previous literature has established the existence of a striking sexual dimorphism in the rodent LC. Compared with males, the female LC contains more NA neurons with higher dendritic complexity and length and LC cells in females respond more robustly to stress (Pinos et al., 2001; Curtis et al., 2005; Bangasser et al., 2011; Guajardo et al., 2017; Mulvey et al., 2018).

Furthermore, a rabies virus tracing study reported sex differences in the pattern of input connectivity onto LC-NA neurons (Sun et al. 2020). Here, we addressed whether there are sex differences in afferent input connections onto functionally/projection identified LC-NA subpopulations of neurons. Interestingly, sex differences were only apparent in inputs to amygdala projecting LC-NA neurons. These inputs were from the hippocampus, the hypothalamic posterior nucleus, the pedunculopontine nucleus and the endopiriform area. These are different input regions compared with the Sun et al. paper which examined inputs to general (not defined by function or projection) LC-NA neurons, suggesting that sex related differences in afferent inputs to LC-NA neurons vary across functionally defined LC neuronal subclasses. In female mice, amygdala-projecting LC neurons received preferential inputs from the hippocampal region. Both the hippocampus and the LC-NA system participate in learning and memory, contextual processing and fear discrimination (Wagatsuma et al., 2017; Kaufman et al., 2020; Seo et al., 2021; see reviews, Hagena et al., 2016; Giustino & Maren, 2018). Furthermore, maladaptive aversive context generalization is a key feature of trauma-related disorders such as PTSD, a condition which is more prevalent in females (Kessler et al., 1995, 2012; Kindt, 2014). In future studies, it will be important to examine the contribution of this hippocampus-LC-amygdala circuit in aversive generalization.

Despite its small size and limited number of neurons, the LC-NA system exerts an outsized influence on brain function and is an important target for the treatment of neurodegenerative and psychiatric disorders (Weinshenker, 2018; Morris et al., 2020; Poe et al., 2020; Breton-Provencher et al., 2021). Emerging evidence suggests that the LC is modularly organized and can exhibit context dependent neural processing. The findings of our study propose common gross-level and specific fine-level afferents as a potential explanation for cell type specific functionality and context dependent modular processing in LC. It will be important in future studies to determine whether and how different biased inputs regulate distinct LC-NA subpopulations during behavior. Importantly, in addition to differences in afferent inputs, other factors may also contribute to LC’s modular functionality including cell type specific variations in local circuit connectivity or modulatory receptor expression as well as differences in intrinsic excitability, release regulation and uptake and/or adrenergic receptor expression in target structures. Future work aimed at further understanding of how differences in afferents interact with other features of LC organization and regulation will be invaluable in understanding this elusive brain region and provide new treatment targets for brain diseases.

## Materials and Methods

### Animals

Nine- to twelve-week-old male and female C57BL/6 or noradrenaline transporter-Cre (NAT-cre) mice (CB57BL/6 background) were used in all experiments. NAT-cre mice were provided by Dr. Thomas McHugh. The vivarium was maintained at a constant temperature (23 ± 1 °C) with a 12:12 light:dark cycle (lights on 7 a.m. to 7 p.m.). Food and water were provided *ad libitum*. All experiments were approved by the Animal Care and Use Committees of the Riken Center for Brain Science.

### Viruses and surgery

Plasmid for adeno-associated virus-CAG promoter-flip-excision-flippase (FLP) recognition site-TVA receptor fused to mCherry (pAAV-CAG-FLEX(FRT)-TC, titer estimated to be 1.893 and 9.56 x 10^12^, Schwarz et al., 2015) was a gift from Dr. Liqun Luo (Addgene plasmid #67827). Plasmid for AAV-FRT site-human histone 2B-human influenza hemaggluttin (HA)-2A self-cleaving peptide-glicoprotein N2c strain (pAAV-CBA-fDIO-H2B-3xHA-P2A-N2cG, titer estimated to be 5.28 × 10^12^ and 1.09 × 10^13^) was produced and package in our lab. Plasmid for the envelope glycoprotein of subgroup A-pseudotyped CVS-N2cΔG fused with EGFP (EnvA-CVS-N2cΔG-EGFP, titer estimated to be 2 × 10^7^, Reardon et al., 2016) was a gift from Dr. Thomas Jessell (Addgene plasmid #73461, Reardon et al., 2016). We thank Dr. Andrew Murray for gifting the Neuro2A cell lines for packaging (Reardon et al., 2016). Canine adenovirus (CAV)-FLEX-Flp (titer estimated to be 2.8 × 10^12^) was obtained from the Montpellier vector core. Retrograde AAV (serotype 2)-FLEx-codon-optimized FLP Recombinase (FlpO) (titer estimated to be 2.03 × 10^12^ and 1.09 × 10^13^) was produced and packaged in our lab.

Mice were anesthetized with isoflurane (3% for induction, 1-1.5% for maintenance) during stereotaxic procedures. For LC subpopulations analysis, 0.3 μl of green or red Retrobeads (Lumafluor Inc.) injections were done into ipsilateral LA/B (from bregma: anterior/posterior (AP): −1.70 mm, medial/lateral (ML): +3.4 mm, dorsal/ventral (DV): −4.3 mm) and IL (from bregma: AP: +1.70 mm, ML: 0.4 mm, DV: −2 mm). Mice were sacrificed for analysis 1 week after surgery.

For cell-type specific trans-synaptic rabies tracing, 0.4 μl of CAV-FLEX-flp and AAV2retro-CBA-FLEX-flpO (mixture 1:1) were injected into LA/B or IL and 0.6 μl of AAV9/2-CAG-FLEX(FRT)-TC and AAV9/2-CBA-fDIO-H2B-3xHA-P2A-N2c(G) (mixture 1:1) into LC (from bregma: AP: −5.4 mm, ML: 0.9 mm, DV: −3.65 mm). Combinations of these viruses were used to reduce individual viral tropism, something we’ve observed in other neural circuits. After 3-4 weeks, mice received injections of 0.6 μl of EnvA-CVS-N2cΔG-EGFP into LC. Ten days after the second surgery, mice were sacrificed for analysis. Viral injection volumes were based on previous publications procedures (Schwarz et al., 2015; Soya et al., 2017; Breton-Provencher & Sur, 2019).

### Histology and immunohistochemistry

For tissue collection, mice were overdosed with isoflurane and perfused with 4% paraformaldehyde in PBS. After post fixation, the brains were sliced using a cryostat. For subpopulation analysis in LC, 50 μm coronal sections containing LA/B, IL and LC were obtained. For retrograde tracing experiments (Fig. 1), LC-containing sections were washed in 0.1 M PBS with 0.3% Triton-X 100 (PBST), followed by 30 min incubation with Donkey Serum 2% in PBST at room temperature. The primary antibody used was sheep anti-tyrosine hydroxylase (TH) (1:2000, AB1542, Chemicon). After overnight incubation at 4 °C, sections were washed in PBS three times and incubated in a PBST solution containing the secondary antibody (Alexa Fluor 647-conjugated donkey anti-sheep, 1:1000, A21448, Invitrogen) and nuclear dye, Hoechst (1:5000-1:10000, H21492, Invitrogen). Following PBS washes at room temperature, sections were mounted with ProLong Gold antifade reagent (P36934, Invitrogen) onto slides and cover slipped. LA/B- and IL-containing coronal sections were mounted right after sectioning to verify retrograde tracers’ expression.

For cell-type specific rabies trans-synaptic tracing, 50 μm coronal sections were collected throughout the entire brain (from bregma AP: ~ +3 mm to AP: ~ −7.20 mm) and every 3^rd^ section was used for immunostaining. For starter cell identification immunohistochemistry in LC-containing sections, primary antibodies used were mouse anti-HA (1:2000, M180-6, MBL), rabbit anti-GFP (1:5000, A11122, Invitrogen) and sheep anti-TH (1:2000, AB1542, Chemicon). The secondary antibodies used were Alexa Fluor 488-conjugated donkey anti-mouse (1:1000, A21202, Invitrogen), Alexa Fluor 594-conjugated donkey anti-rabbit (1:1000, A21207, Invitrogen) and Alexa Fluor 647-conjugated donkey anti-sheep (1:1000, A21448, Invitrogen) and nuclear dye, Hoechst (1:10000, H21492, Invitrogen).

For whole-brain inputs analysis on sections which did not contain LC, the primary antibody used was rabbit anti-GFP (1:5000, A11122, Invitrogen) and the secondary antibody used was Alexa Fluor 488-conjugated donkey anti-rabbit (1:1000, A21202, Invitrogen). In addition, we performed nuclear staining using Hoechst dye (1:10000, H21492, Invitrogen).

### Microscopy

For retrograde tracing experiments (Fig. 1), LC, LA/B and IL images were obtained using a confocal laser scanning microscope (FluoView FV-1000, Olympus) with a 20X (in LA/B and IL sections) or 40X (in LC sections) objective. For starter cell identification in rabies tracing experiments, LC images were obtained using a confocal laser scanning microscope (FluoView FV-1000, Olympus, Japan) with a 40X objective. For whole-brain input analysis on sections that did not contain LC, sections were obtained using a digital slide scanner (NanoZoomer s60, Hamamatsu Photonics, Japan) with a 20X objective. All images obtained from confocal and NanoZoomer analysis were taken using tiling and z-stack functions. To identify the correct injection sites of retrograde viruses in LA/B and IL, cannula tracks were reconstructed under a fluorescence microscope.

### Image registration and signal identification

For the rabies virus input quantification, the registration process of the whole-brain sections was based on previous publications (Pollak Dorocic et al., 2014; Zhang et al., 2016; Sun et al., 2020). Every 3^rd^ section was processed for analysis. Sections containing the rabies virus injection and neighboring regions (approximately from bregma −5.04 to −5.80) were excluded from the inputs analysis following previous LC rabies virus protocols (Schwarz et al., 2015; Do et al., 2016; Sun et al., 2020). Coronal images of individual sections were cropped from NanoZoomer images using image viewing software (NDP.view2, Hamamatsu, Japan). Image stacking and alignment of left and right hemispheres were done using Fiji ImageJ (National Institutes of Health, USA, (Schindelin et al., 2012). For matching procedures, Mouse Brain Reference Atlas (Allen Institute, 2011) online templates were overlapped to individual sections using the Big Warp plugin of Fiji ImageJ. This plugin allows deformation of reference templates onto individual sections to match landmarks. Identification and quantification of input cells throughout the brain was conducted manually using the Cell Counter plugin of Fiji ImageJ. Cells located between regions were assigned to the nearest region. To control for variations in the number of input cells across animals, absolute input numbers per region were normalized by the total number of inputs from all brain regions.

Animals without starter cells (helper -cytoplasmic expression of mCherry for TVA and nuclear expression of Alexa Fluor 594 for HA- and rabies-EGFP expressing) in LC were excluded from analysis.

### Brain region annotation and data analysis

Whole-brain input annotation was done according to the Allen Mouse Brain Reference Atlas region classification. This classification is represented in a tree-like hierarchical structure. The broadest level of hierarchy or “root” is the gray matter, fiber tracts and ventricular systems which are then further subdivided into “leaves” which represent the divisions of the root level. Nodes are branch points in the hierarchical structure giving rise to leaves (e.g., gray matter is a node for cerebrum, brainstem and cerebellum leaves). Leaves can then be “nodes” for finer subdivisions and so forth (Sunkin et al., 2013; Wang et al., 2020 and Takata et al 2021; see Allen Institute). We considered the “root” as the first “node”. Levels of organization used for analysis were similar to those used by Sun et al., 2020: gross (mainly 4^th^ “node” regions), moderate (mainly the 5^th^ “node” subregions, subdivisions of the previous level) and fine (mainly structures belonging to the 6^th^-7^th^ “nodes”). Statistical analysis was done using GraphPad Prism 6.0 or 9.4 (GraphPad Software, Inc., USA). Data was presented as mean ± s.e.m. Normality and homogeneity of variances analysis were conducted to decide appropriate statistical tests. To evaluate significance of fraction of inputs, Mann-Whitney U tests were performed, and significance level was set at P<0.05. Statistical analyses were based on previous publications (Ährlund-Richter et al., 2019; Sun et al., 2020).

## Acknowledgements

we thank Yuri Ishizu and Nisha Ponneduthamkuzhy for technical support and members of the Johansen lab for improving the manuscript with their insightful comments.

