## Supplementary Figures 1-4 for "Whole-brain afferent input mapping to functionally distinct brainstem noradrenaline cell types"

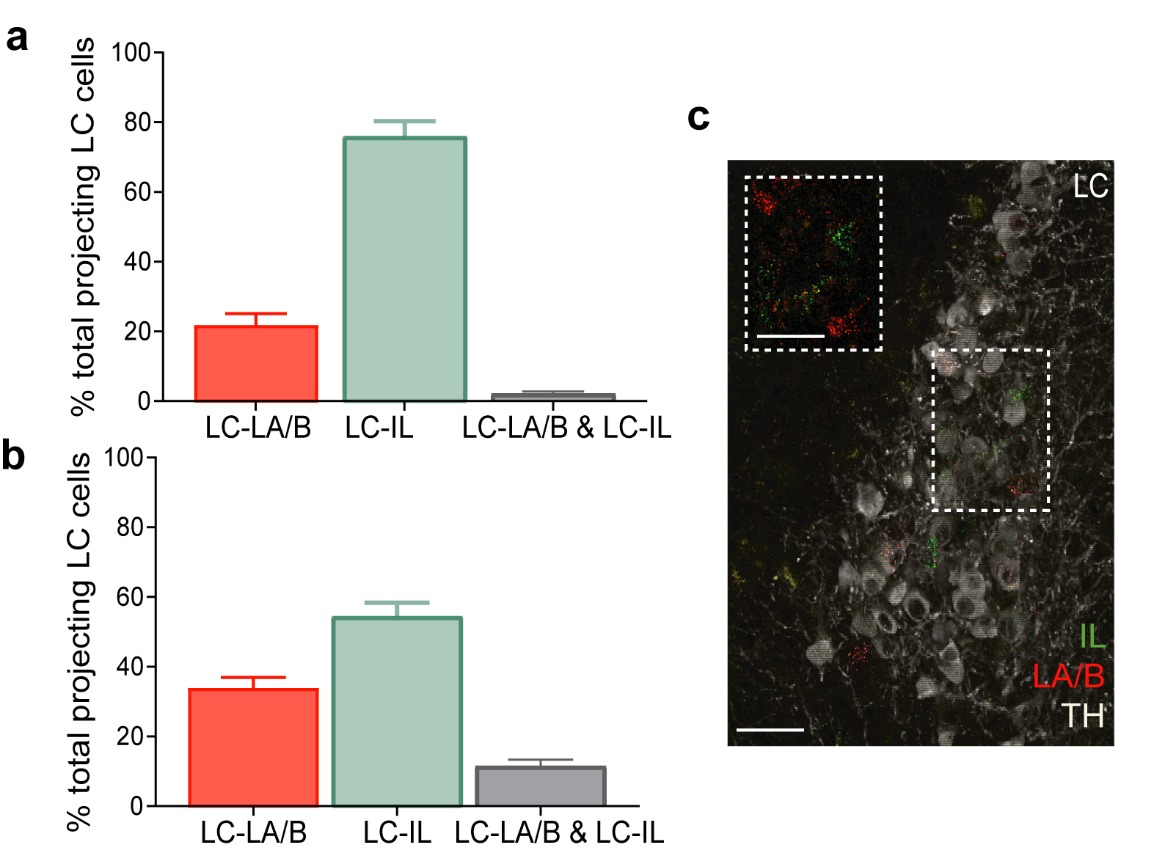
Supplementary Figures

***Supplementary Figure 1. Locus coeruleus (LC) projections to lateral and basal nuclei of the amygdala (LA/B) and infralimbic cortex (IL) in male and female mice.***

***a)*** *Percentage of total LC cells to LA/B, IL or both regions in male mice (n= 2, mean, s.e.m.).* ***b)*** *Percentage of total LC cells to LA/B, IL or both regions in female mice (n= 2).* ***c)*** *Example of female LC. Inset: close-up example of input cells from LC to LA/B (red) and IL (green). Tyrosine hydroxylase (TH) immunostaining to label LC-NA neurons pseudocolored in white. Scale: 50 μm.*

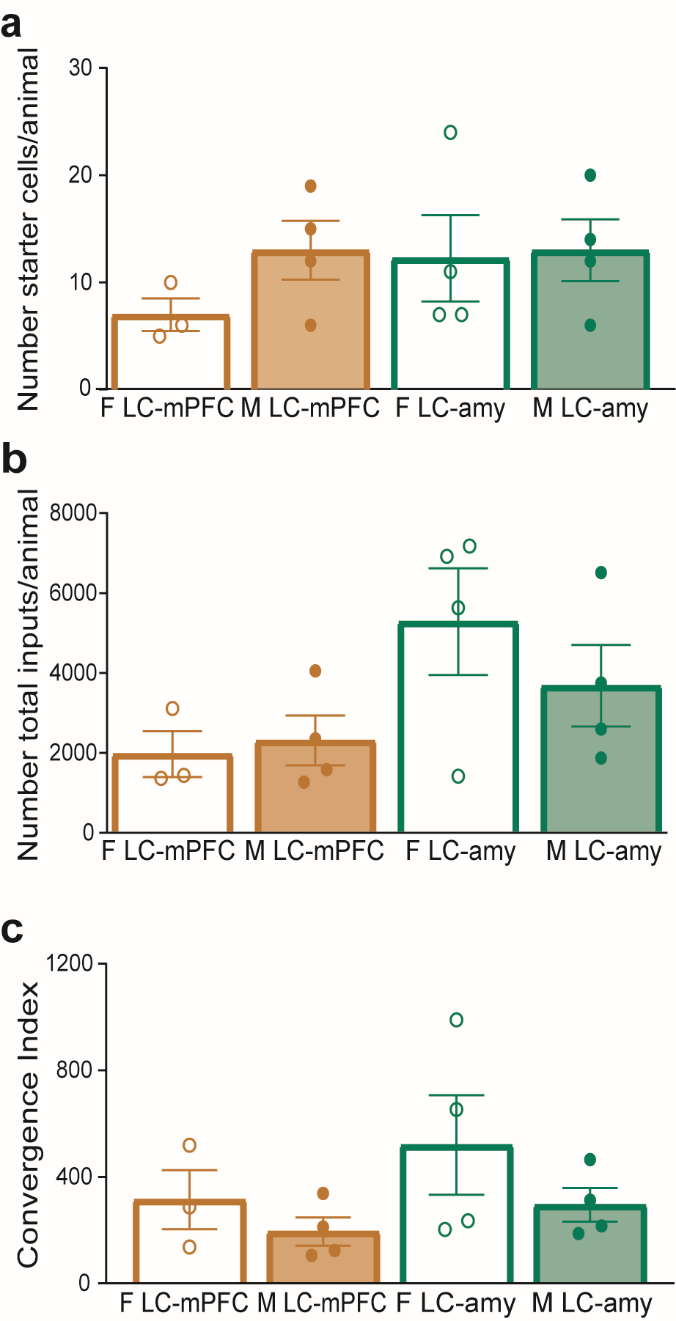

***Supplementary Figure 2. Sex-specific quantification for rabies virus experiments***

***a)*** *Number of starter cells for female (F) and male (M) LC-amydala (LC-amy) and LC-mPFC groups.* ***b)*** *Number of total inputs for female and male LC-amy and LC-mPFC groups.* ***c)*** *Convergence Index for female and male LC-amy and LC-mPFC groups. Circles represent individual animal values (fem LC-amy, n= 3; male LC-amy, n= 4; fem LC-mPFC, n= 4; male LC-mPFC, n= 4; mean ± s.e.m.).*

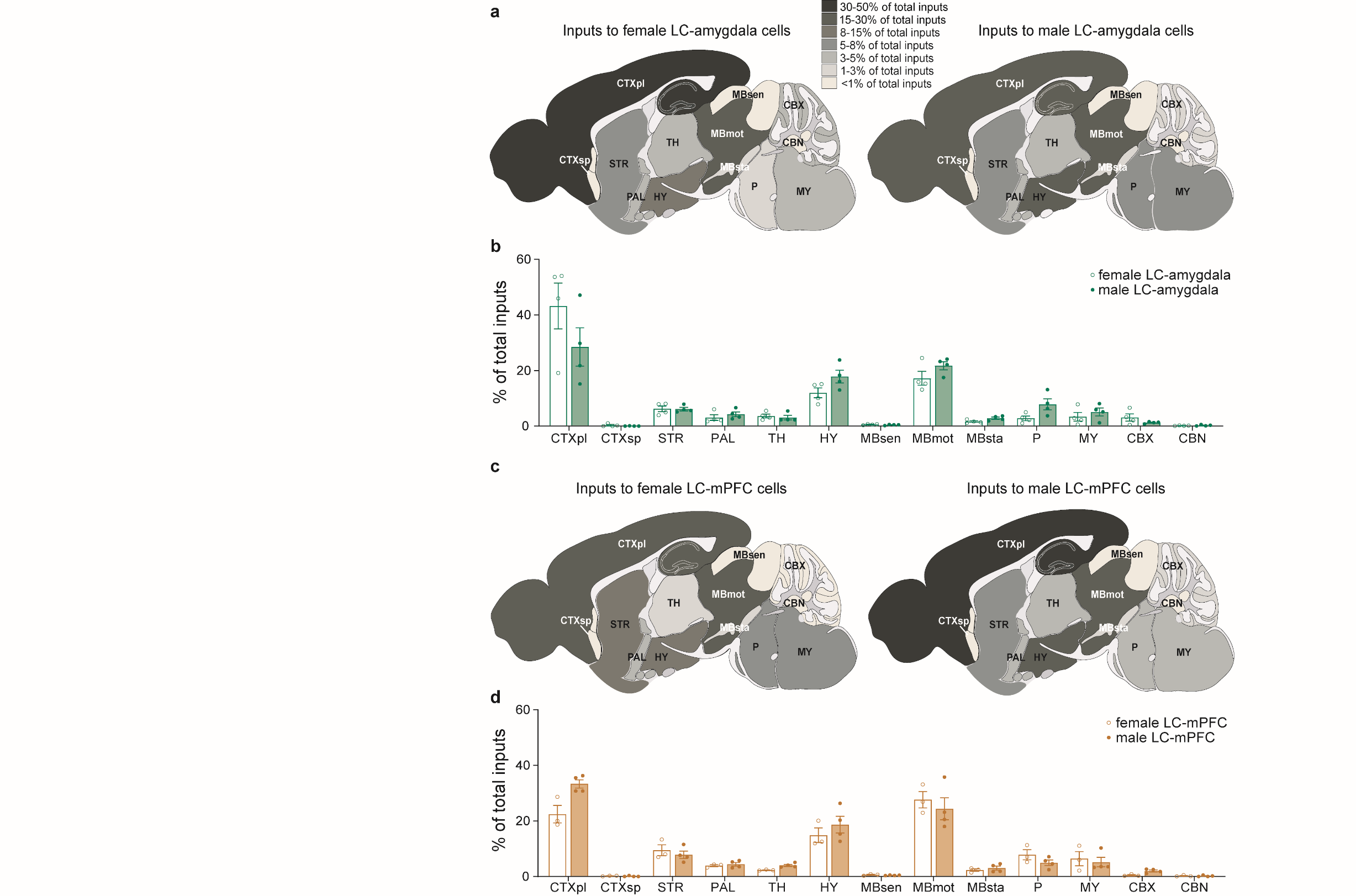

***Supplementary Figure 3. Gross level of afferent input organization to amygdala and mPFC projecting LC-NA neurons in female and male mice.***

***a)*** *Whole-brain input distribution. Left: colorized schematic illustrating proportion of inputs to female amygdala-projecting LC-NA cells. Right: colorized schematic illustrating proportion of inputs to male amygdala-projecting LC-NA cells.* ***b)*** *Percentage of total inputs from the whole brain to female and male LC-amygdala groups (LC-amygdala group: female, n= 3; male, n= 4).* ***c)*** *Whole-brain input distribution. Left: proportion of inputs to female mPFC-projecting cells. Right: proportion of inputs to mPFC-projecting cells.* ***d)*** *Percentage of total inputs from the whole-brain to female and male LC-mPFC groups (LC-mPFC group: female, n= 4; male, n= 4). Circles denote individual animal values. CTXpl: cortex, plate. CTXsp: cortex, subplate. STR: striatum. PAL: pallidum. TH: thalamus. HY: hypothalamus. MBsen: midbrain, sensory-related. MBmot: midbrain, motor related. MBsta: midbrain, behavioral state related. P: pons. MY: medulla. CBX: cerebellar cortex. CBN: cerebellar nuclei.*

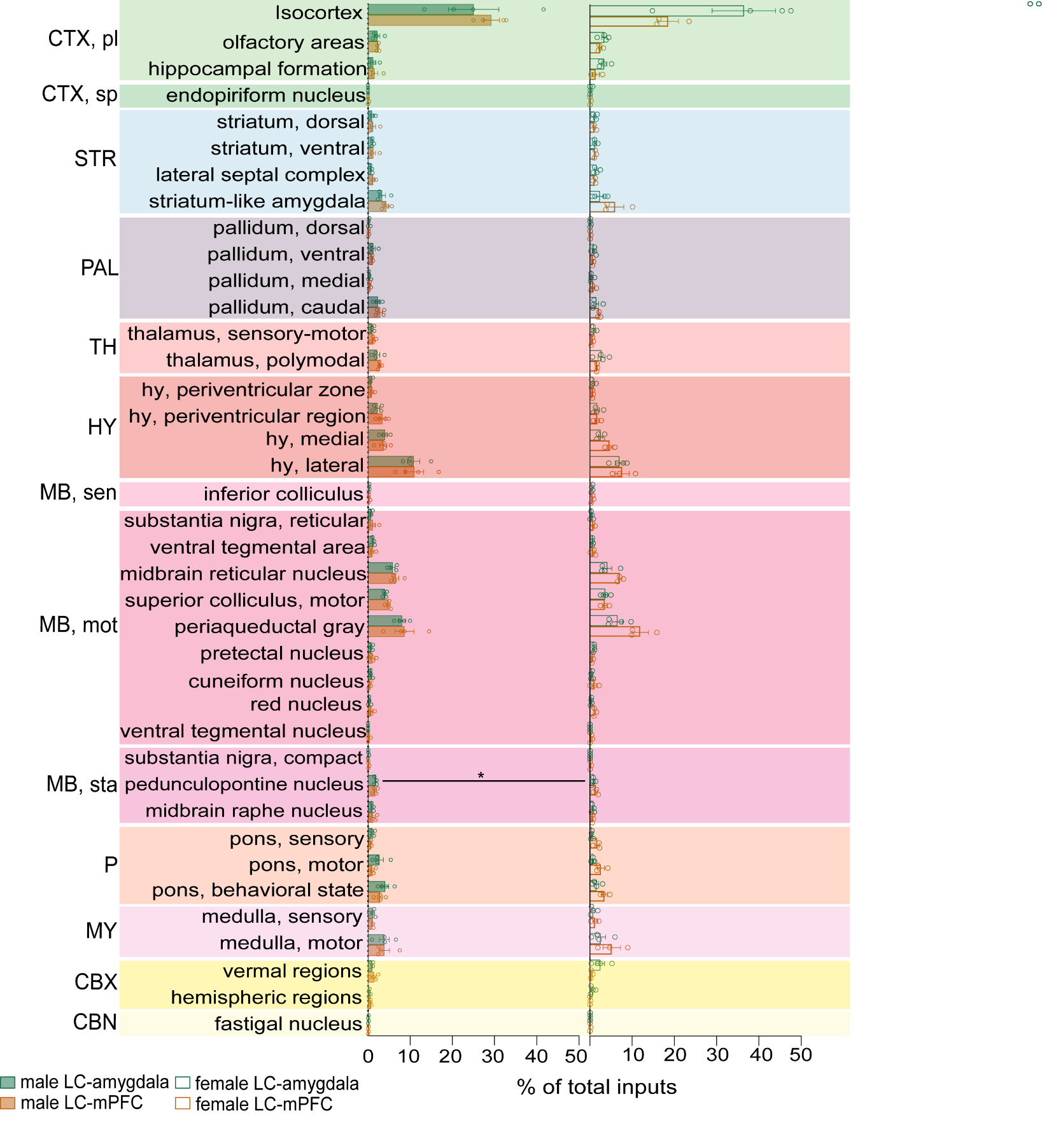

***Supplementary Figure 4. Moderate level of input organization in female and male mice****.*

*Percentage of total inputs the moderate level of organization regions male mice. *P < 0.05. Circles represent individual animal values.*
